# Bacterial diversity in deep-sea sediments under influence of asphalt seep at the São Paulo Plateau

**DOI:** 10.1101/753616

**Authors:** Luciano Lopes Queiroz, Amanda Gonçalves Bendia, Rubens Tadeu Delgado Duarte, Diego Assis das Graças, Artur Luiz da Costa da Silva, Cristina Rossi Nakayama, Paulo Yukio Sumida, Andre O. S. Lima, Yuriko Nagano, Katsunori Fujikura, Hiroshi Kitazato, Vivian Helena Pellizari

## Abstract

Here we investigated the diversity of bacterial communities from deep-sea surface sediments under influence of asphalt seeps at the Sao Paulo Plateau using next-generation sequencing (NGS) method. Sampling was performed at North São Paulo Plateau using the human occupied vehicle Shinkai 6500 and her support vessel Yokosuka. The microbial diversity was studied at two surficial sediment layers (0-1 and 1-4 cm) of five samples collected in cores in water depths ranging from 2,456-2,728 m. Bacterial communities were studied through sequencing of 16S rRNA gene on the Ion Torrent platform and clustered in operational taxonomic units. We observed high diversity of bacterial sediment communities as previously described by other studies. When we considered community composition, the most abundant classes were Alphaprotebacteria (27.7%), Acidimicrobiia (20%), Gammaproteobacteria (11.3%) and Deltaproteobacteria (6.6%). Most abundant OTUs at family level were from two uncultured bacteria from Actinomarinales (5.95%) and Kiloniellaceae (3.17%). The unexpected high abundance of Alphaproteobacteria and Acidimicrobiia in our deep-sea microbial communities may be related to the presence of asphalt seep at North São Paulo Plateau, since these bacterial classes contain bacteria that possess the capability of metabolizing hydrocarbon compounds.

## Introduction

Deep-sea ecosystems represent most of Earth surface. The seabed is composed of several types of habitat as hydrothermal vents, cold seeps, seafloor and subseafloor (Jørgensen and Boetius 2007; Orcutt et al. 2011). Seafloor sediments are particularly interesting due to their geochemical characteristics, sedimentary dynamics and greater habitat stability that are important factors to structuring communities of macro and microorganisms. Research on microbial diversity in superficial sediment habitat has been intensified, in an effort to better understanding how spatial and temporal patterns are determined (Zinger et al. 2011; Nemergut et al. 2013).

Spatial distribution of microorganisms in deep-sea habitats has been studied in several locations, from Arctic sediments in the Pacific Ocean (Li et al. 2009), to Siberian continental margin (Bienhold et al. 2012), eastern South Atlantic sediments near Angolan coast (Schauer et al. 2010) and Southwestern Atlantic pockmarks close to the Brazilian coast (Giongo et al. 2015). Although there are few studies in deep-sea habitats from South Atlantic Ocean, the knowledge of how, what and where microorganisms inhabit is incipient compared with similar environments from North Atlantic or other better studied deep-sea basins.

The Brazilian coast is known for the presence of large oil fields under seafloor sediments (Coward et al. 1999; Winter et al. 2007). Campos and Santos basins are important oil-producing areas of Brazil, responsible for more than 71% of the country’s oil production (Almada and Bernardino 2017). Considering the existence of oil and gas reservoirs in these basins, it was expected that chemosynthetic ecosystems exist and a joint Japanese-Brazilian Iatá-Piúna cruise was conducted to investigate that hypothesis. This cooperative project integrated the Quest for the Limits of Life (QUELLE) 2013 carried out by JAMSTEC (Japan Agency for Marine-Earth Science and Technology). During the cruise that explored the deep seafloor of the North São Paulo Plateau in Espírito Santo Basin (2500-3600 m), asphalt seeps were found at a depth of 2,700 m colonised by non-chemosynthetic megafaunal organisms (Fujikura et al. 2017). They also found that, in non-asphalt seeps areas, outcrops of mudstone were covered by black manganese oxide crusts and nodules were also present (Aguiar et al. 2014; Fujikura et al. 2017; Jiang et al. 2018). These two particular conditions of the study area may be important factors determining patterns of bacterial diversity.

A previous study carried out by the Iatá-Piúna consortium (Jiang et al. 2018) using PCR-DGGE method found high and widespread dominance of Proteobacteria and Firmicutes at sediment samples, including asphalt seep area. The two predominant species were *Erythrobacter citreu*s strain VSW309 detected in hydrothermal vents and *Thalossospira xianhensis* strain MT02 a hydrocarbon-degrading marine bacterium. They also found that microbial community composition between sediment core depths was different.

Here we investigate the diversity of bacterial communities from deep-sea surface sediments under the influence of asphalt seeps at the Sao Paulo Plateau using next generation sequencing (NGS). Bacterial community assembly was accessed using high-throughput 16S rRNA gene sequencing on an Ion Torrent PGM platform and by quantitative amplification (qPCR) with the aim of (1) describing bacterial diversity and (2) estimating bacterial populations present in sediment depth layers.

## Material and Methods

### Description of sampling sites

Sediment samples were collected during 2^nd^ leg of ‘*Iatá-piuna cruise*’ expedition, a collaborative project between Brazil and Japan inserted in the QUELLE (Quest for Limit of Life) initiative from JAMSTEC (Japan Agency for Marine-Earth Science and Technology). Sediments samples were collected using the HOV ‘*Shinkai 6500*’ and support vessel ‘*Yokosuka*’ in Sao Paulo Plateau located off the coast of Espírito Santo and Rio de Janeiro states, composing the Campos and Espírito Santo basins.

The study area was the North São Paulo Plateau (Figure 1), this region is located between coordinates 20°30’ – 21°30’ S and 39°30’ – 38°30’ W. A total of 5 sediment cores were sampled by push corers (30 cm in length and 10 cm in diameter) operated by the manipulators of Shinkai 6500. Cores were subsampled on board at depth intervals of 0-1 cm, 1-4 cm, 4-7 cm, 7-10 cm and 10-13 cm. Each subsample was placed in sterile sample bags and stored at -20 °C. The top two layers (0-1 and 1-4 cm) of sampled sediment cores from North Sao Paulo Plateau were selected (Table 1) due the presence of asphalt seep. Samples N11, N12, and N13 were associated with asphalt seep and N06 and N14 were from background deep-sea, distant between them 2 to 5 km.

**Table 1.** Coordinates, depths and alphalt seep presence/absence of sediment samples from North São Paulo Plateau.

| Dive<br>No. | Sample | Core<br>No. | Latitude (S) | Longitude<br>(W) | Seafloor<br>depth (m) | Asphalt<br>seep area | Sediment Layer |
| --- | --- | --- | --- | --- | --- | --- | --- |
| 1343 | 06 | 02 | 20°41'37.57" | 38°38'11.86" | 2728 | No | N06-1cm<br>N06-4cm |
| 1345 | 11 | 05 | 20°43'8.05" | 38°39'6.23" | 2720 | Yes | N11-1cm<br>N11-4cm |
| 1346 | 12 | 03 | 20°43'54.20" | 38°39'43.76" | 2651 | Yes | N12-1cm<br>N12-4cm |
| 1346 | 13 | 04 | 20°43'54.20" | 38°39'43.76" | 2651 | Yes | N13-1cm<br>N13-4cm |
| 1347 | 14 | 03 | 20°44'17.20" | 38°40'7.61" | 2456 | No | N14-1cm<br>N14-4cm |

**Figure 1.**
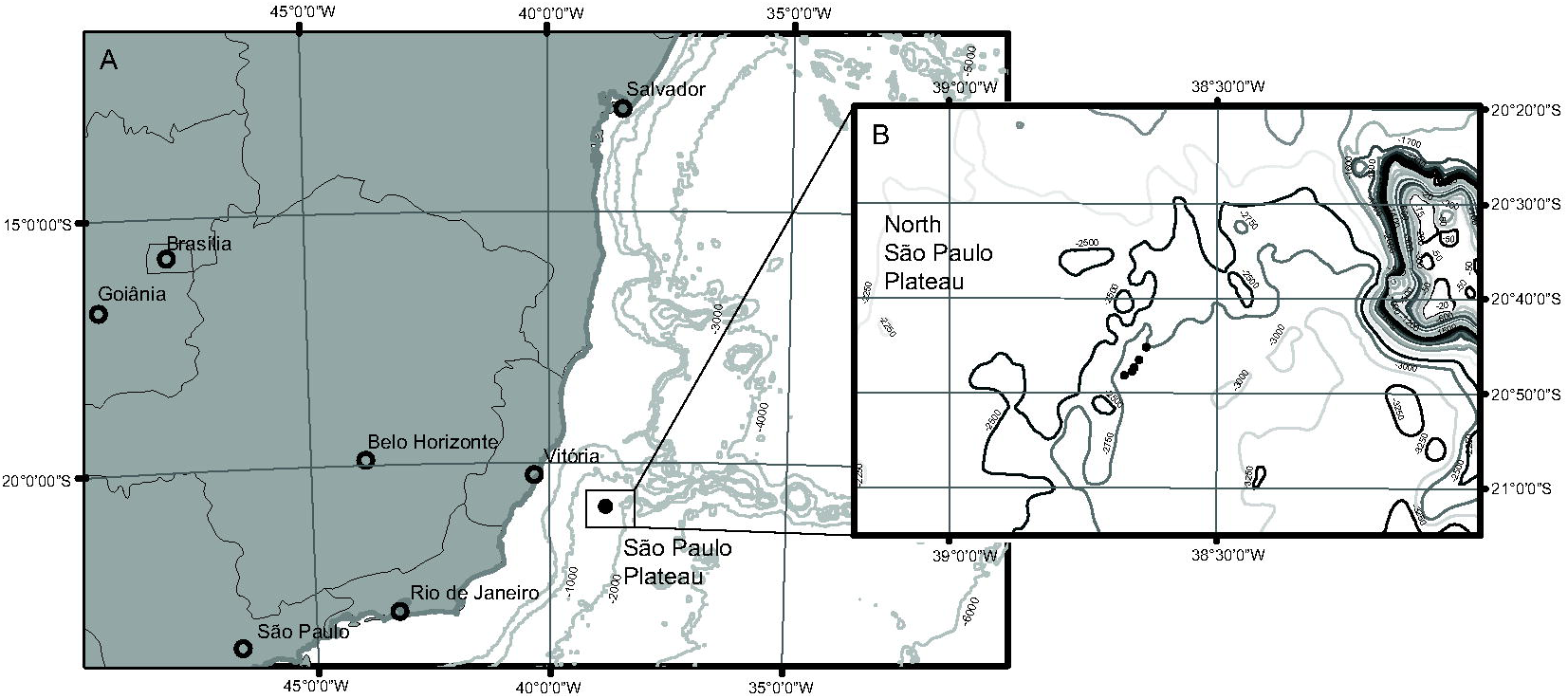
Map and location of samples in the São Paulo Plateau. (A) Location São Paulo Plateau and (B) distribution of superficial sediments samples in North São Paulo Plateau.

### DNA extraction

Total community DNA was extracted from the top layers (0-1 and 1-4 cm) of five sediment cores from North Sao Paulo Plateau (Table 1) using PowerSoil® DNA Isolation Kit (MO BIO Laboratories Inc., Carlsbad, CA, USA) following manufacturer’s instructions with adaptations: sample consisted of 0.5 g of homogenised sediment and after mechanic cell lysis, a thermal shock step was added, heating samples at 55 °C for five minutes followed by 1 minute at -20 °C.

The integrity of extracted DNA was evaluated by electrophoresis in 1% agarose gel with TAE 1X (Tris 0.04M, glacial acetic acid 1M, EDTA 50 mM; in pH 8), visualised with SYBR^®^Safe (Invitrogen, Paisley, UK), and Lambda Hind III (Life Technologies, Carlbad, CA., EUA) used as molecular marker. DNA was quantified using NanoDrop ND-1000 spectrophotometer (Thermo Scientific, Waltham, MA, EUA) and Qubit® dsDNA HS (High Sensitivity) Assay (Life Technologies).

### Ion Torrent PGM Sequencing

The bacterial 16S rRNA gene V3 and V4 variable regions were amplified with primers 341F (5’-CCTACGGGNGGCWGCAG-3’) and 805R (5’-GACTACHVGGGTATCTAATCC-3’) (Herlemann et al. 2011). PCR mixtures contained 0.5 µM of each primer, 0.7 U of Taq DNA Polymerase (Life Technologies, Carlbad, CA., EUA), 1X Buffer, 4 mM of MgCl_2_, 0.2 mM of each dNTP, 0.3 mg/mL BSA (Bovine Serum Albumin) and 4 ng of DNA template. Cycling conditions consisted of 5 min initial denaturation at 95 °C; 2 cycles of 1 min denaturation at 95 °C, 1 min annealing at 48 °C and 1 min extension at 72 °C; 2 cycles of 1 min at 95 °C, 1 min at 50 °C and 1 min at 72 °C; 2 cycles of 1 min at 95 °C, 1 min at 52 °C and 1 min at 72 °C; and 22 cycles of 1 min at 95 °C, 1 min at 54 °C and 1 min at 72 °C. The first few cycles with increasing annealing temperature is an adaptation to avoid mixed-template PCRs bias in the final products (Ishii and Fukui 2001).

Amplicons libraries obtained were purified before emulsion step with Purelink PCR Purification Kit (Life Technologies, Carlbad, CA., EUA) and quantified using Qubit® dsDNA HS (High Sensitivity) Assay (Life Technologies). Emulsion PCR was carried out using Ion OneTouch 2™ Instrument, using the Ion PGM™ Template OT2 Reagents 400 Kit and enriched with OneTouch ES (Life Technologies). The sequencing of libraries was carried out in an Ion PGM™ System, using the Ion PGM Sequencing 400 Kit and deposited in two Ion 318 chip Kit v2 following the manufacturer’s protocol (Life Technologies).

### Quantitative PCR (qPCR)

Enumeration of bacterial populations was carried out by qPCR, performed in triplicates using SYBR Green I system detection (Invitrogen). Previous to qPCR, DNA was purified with the OneStep™ PCR Inhibitor Removal Kit (Zymo Research, USA) and diluted 1:5. The bacterial primers used were 27F 5’-AGAGTTTGATCMTGGCTCAG-3’ and 518R 5’-GTATTACCGCGGCTGCTGG-3’ (Muyzer et al. 1993). Each reaction contained 12.5 µL of Platinum^®^ Quantitative PCR SuperMix-UDG (Invitrogen), 0.2 µM of each primer, 0.5 µL of BSA (Bovine Serum Albumin), 5 µL of template DNA and ultra-pure water to complete 25 µL final volume. Amplification conditions to bacterial primers were: initial denaturation at 95 °C for 10 min, followed by 40 cycles of denaturation at 95 °C for 1 min, annealing at 56 °C for 30 seconds and elongation at 72 °C for 30 seconds. The specificity of the reaction was verified against the denaturing curve with temperatures ranging from 72 °C to 96 °C. Data were analysed using Applied Biosystems software, and values of cycle threshold (Ct), logarithmic correlation (R^2^) between number of cycles and DNA quantity in samples and reaction efficiency were calculated. As positive controls of qPCR reactions, serial dilutions of 16S rRNA PCR product of *Escherichia coli* amplified with the primers 27F-1401R (Lane 1991) were used. Thus, values of Ct obtained in each reaction were utilised to determine the absolute quantity of DNA in samples and result were represented by the 16S rRNA gene copy numbers per gram of sediment.

### Bioinformatics

The first filter step was carried out using PGM software to remove low quality and polyclonal sequences. We performed bioinformatics analysis using the Brazilian Microbiome Project (BMP) pipeline (Pylro et al. 2014). BMP pipeline is a combination of VSEARCH (Rognes et al. 2016) and QIIME (version 1.9.0) (Caporaso et al. 2010) software. Using VSEARCH barcodes and primer sequences we removed from *fastq* file, sequences were filtered by length (fastq_trunclen 200) and quality (fastq_maxee 1.0), sorted by abundance and removed singletons. After that, OTUs were clustered and chimeras were removed. We assigned taxonomy using *uclust* method in QIIME and SILVA 16S Database (version n132) as reference sequences (Quast et al. 2013). The OTU table file was converted to BIOM and taxonomy metadata was added. Diversity indices of Chao1, Shannon (log base 2) and Simpson were calculated among samples.

### Statistical analyses

Alpha-diversity analysis were compared between alphalt and non-asphalt seep areas using a *t-*test (Sokal and Rohlf 1995). The 50 most abundant OTUs were filtered and a heat map was constructed considering taxonomic classification (Class and Order) and abundance using Ward’s hierarchical clustering method (ward.d2) (Murtagh and Legendre 2014). The estimated number of bacterial 16S rRNA gene copies were compared between sediment layers by Wilcoxon-Mann Whitney test (Fay and Proschan 2010). Moreover, we performed beta diversity analyses to compare similarities between samples through Principal Coordinates Analysis (PCoA) and using distance matrix of Bray-Curtis, Jaccard, Unweighted and Weighted Unifrac. The differences were tested using Permutational analysis of variance (PERMANOVA) (Anderson 2001). All analyses were carried out using the statistical software R (R Development Core Team 2014), *qiimer, ggplot2* (Wickham 2016), *phyloseq* (McMurdie and Holmes 2013) and *vegan* packages (Oksanen et al. 2013).

## Results

We obtained 520,863 sequences and 5,229 OTUs clustered at 97% of similarity after quality control and bioinformatics analysis from 10 sediment samples using Ion Torrent PGM. The number of sequences varied among samples, ranging from 1,121 sequences in N12-1 to 120,296 in N13-1. Samples N06-2 (3,444), N11-2 (6,566) and N12-1 (1,121) showed a low number of reads, we rarefied all samples to 25,000 reads and excluded those samples from alpha and beta-diversity analysis.

Alpha-diversity analysis showed that the number of observed species (OTU_0.03_) ranged from 904 to 2,282, revealing a wide range of species inhabiting the Sao Paulo Plateau (Table 2). Samples with highest richness indices were N13-1 (2,282), N13-2 (2,184), and N14-1 (2,017). On the other hand, samples with lowest richness indices were N11-1 (904) and N12-4 (1621). Similarly, estimated richness by Chao1 index ranged from 973 to 3072 species. However, in contrast to the higher variability observed in the number of OTUs and estimated richness, the Shannon and Simpson indices were more uniform among samples, ranging from 7.841 to 8.216, and 0.982 to 0.987, respectively (Table 2). We did not find significative differences of alpha-diversity between asphalt and non-asphalt seep areas, except to Simpson index (Suppl. Table 1).

**Table 2.** Richness and alpha diversity data found in North region from São Paulo Plateau and obtained by Ion Torrent;

| Samples | Reads<sup>a</sup> | Observed | Chao1 | Shannon | Simpson |
| --- | --- | --- | --- | --- | --- |
|  |  | species<br>(OTU<sub>0.03</sub>) | (sd) |  |  |
| N06-1cm | 94184 | 1996 | 2587±59 | 8.106 | 0.984 |
| N11-1cm | 38311 | 904 | 973±16 | 7.841 | 0.987 |
| N12-4cm | 65601 | 1702 | 2076±45 | 8.120 | 0.986 |
| N13-1cm | 120296 | 2282 | 3072±70 | 8.216 | 0.985 |
| N13-4cm | 88568 | 2184 | 2888±66 | 8.190 | 0.985 |
| N14-1cm | 81735 | 2017 | 2661±62 | 7.914 | 0.982 |
| N14-4cm | 26579 | 1621 | 1842±30 | 8.043 | 0.983 |

<sup>a</sup> Number of reads of all samples were rarified to 25000 reads before alpha-diversity analysis.

In general, the community composition from all sediment samples was similar, with most sequences classified within the phyla Proteobacteria (45.7%, mean of all samples), Actinobacteria (20.8%), Chloroflexi (3.74%), Acidobacteria (3%), and Gemmatimonadetes (2.1%) (Figure 2). A similar trend was observed when sequences from the phyla Proteobacteria and Actinobacteria were analysed at class level, with Alphaproteobacteria (27.7%), Acidimicrobiia (20%), Gammaproteobacteria (11.3%) and Deltaproteobacteria (6.6%) composing the sediment community (Figure 3).

**Figure 2.**
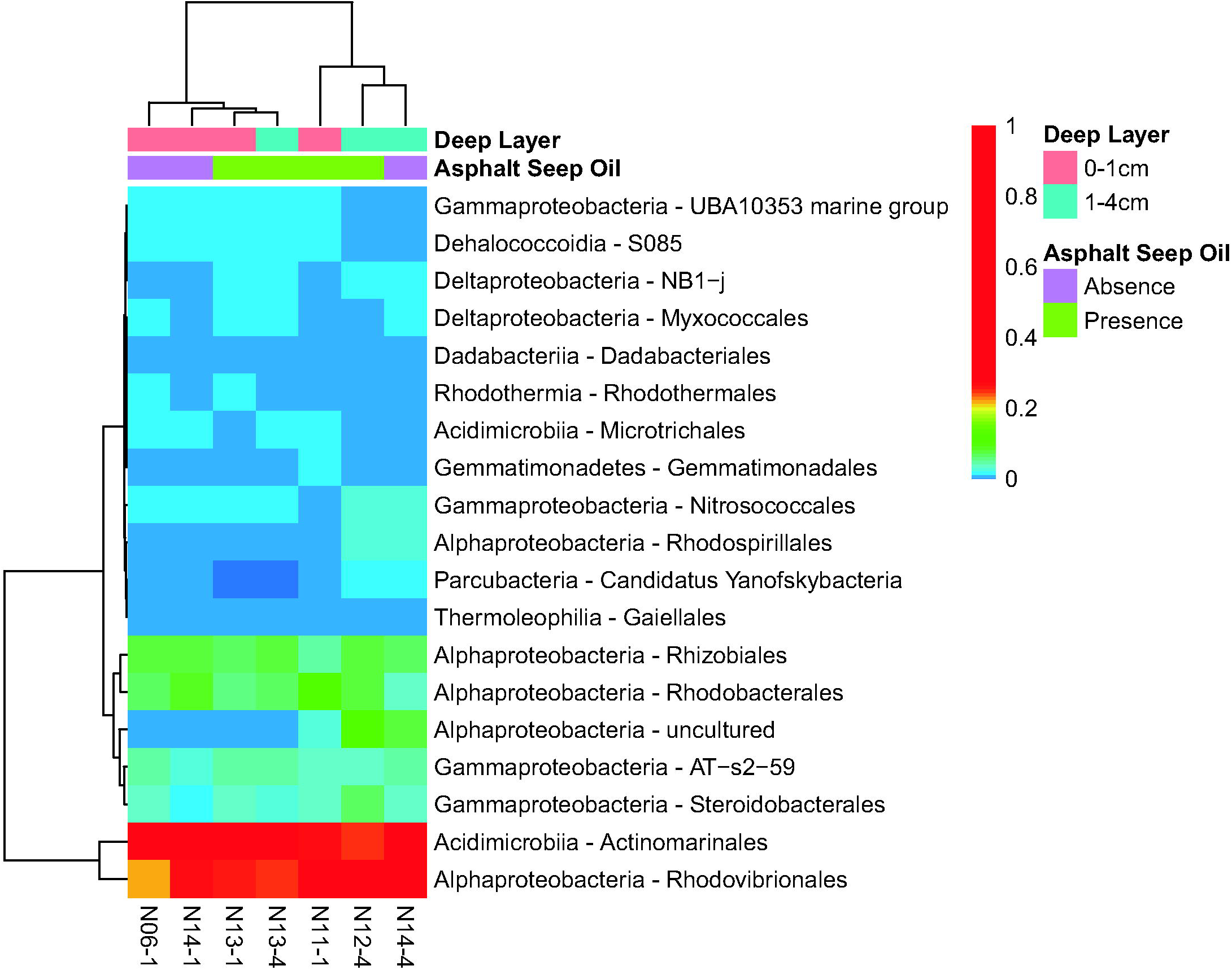
Heat map with 50 most abundant bacterial OTUs (classified by Class and Order) among the different depths and asphalt seep presence/absence in North São Paulo Plateau.

**Figure 3.**
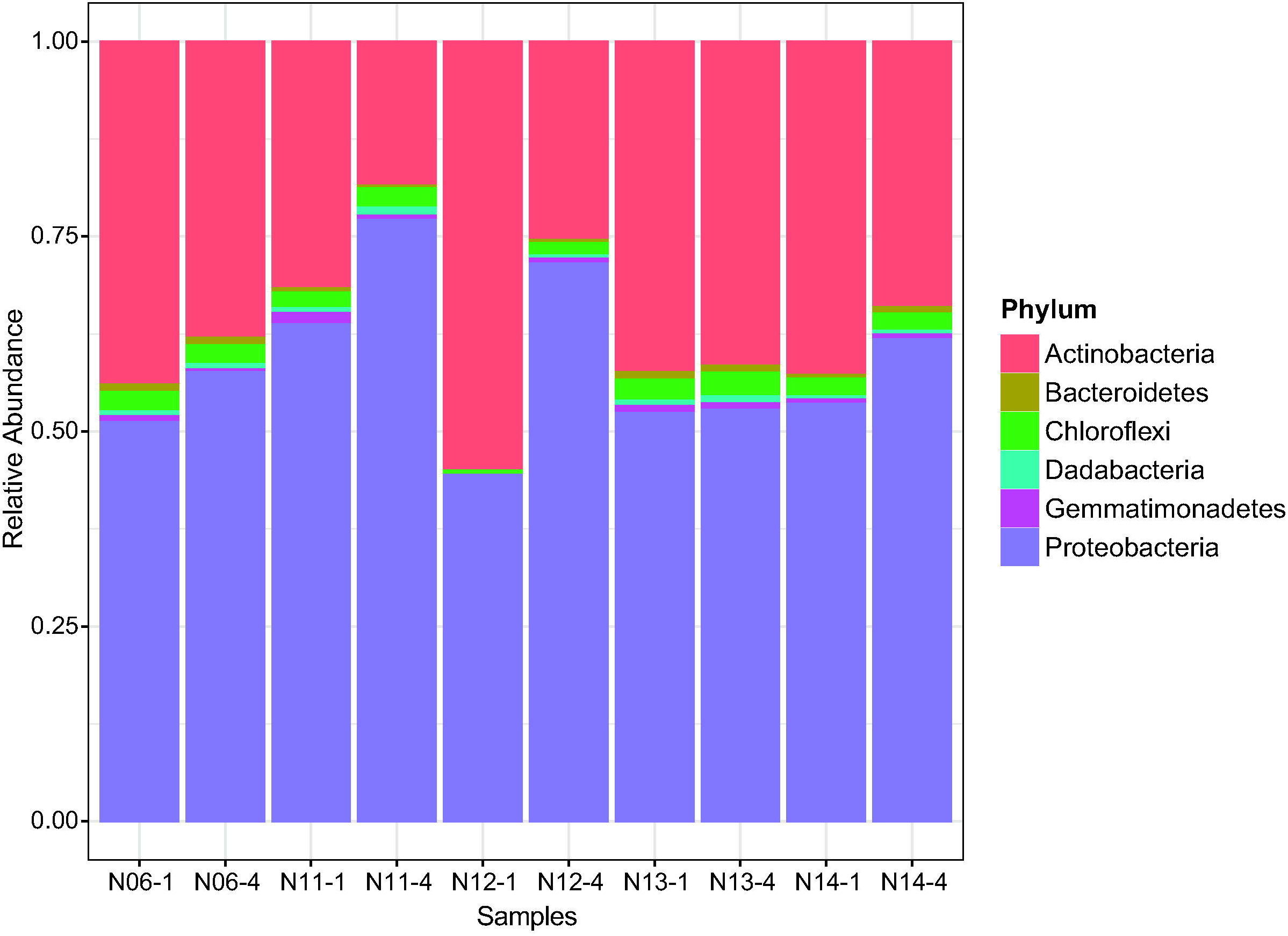
Relative abundance of the most abundant phyla found in North São Paulo Plateau.

The first and thirty most abundant OTUs were an uncultured bacterium of the order Actinomarinales (Acidimicrobiia) (5.95% and 2.75%).Second and forty most abundant OTUs were an uncultured bacterium of the order Rhodovibrionales and family Kiloniellaceae(Alphaproteobacteria) (3.17% and 2.59%), followed by an uncultured bacterium of the order AT-s2-59 (Gammaproteobacteria) (1.91%). Among OTUs classified at genus level, AqS1 (Gammaproteobacteria: Nitrosococcaceae) (0.69%) was found in all samples.

Samples were not clustered by sediment depth or asphalt seep presence/absence in heatmap (Figure 4) and PCoA analyses (Suppl. Figure 1). We did not identify significant correlation between community distance matrix used in the PCoA analysis and samples category (Suppl. Table 2). Heatmap analysis showed the clusterization of two sample groups. Further, OTUs classified at Class and Order were divided in three distinct groups, in which one group was related to Actinomarinales and Rhodovibrionales orders, the second group composed by orders Rhizobiales, Rhodobacterales, AT-s2-59, Steroidobacterales and uncultured Alphaproteobacteria, and a third group formed by less abundant orders.

**Figure 4.**
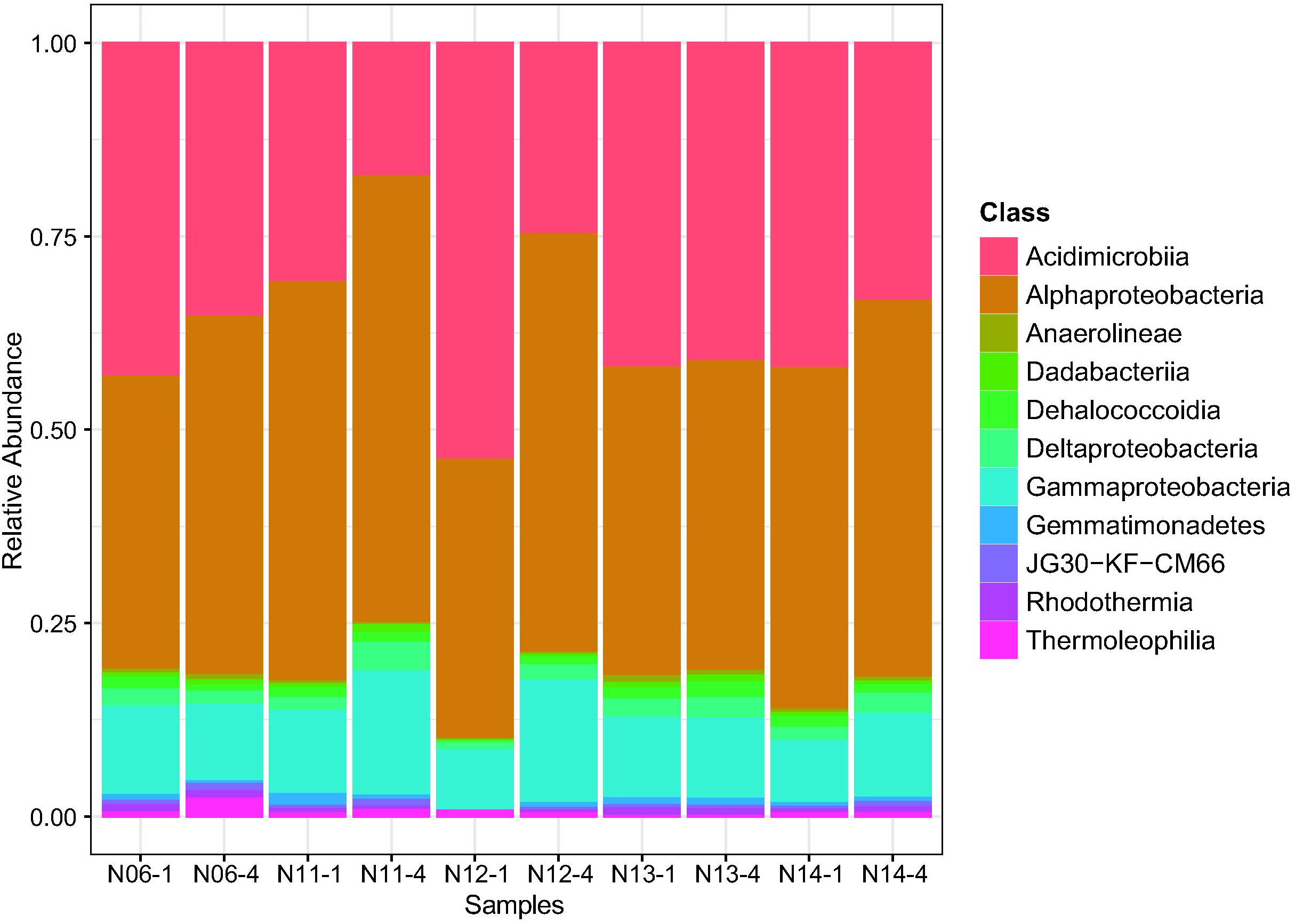
Relative abundance of the most abundant classes found in North São Paulo Plateau.

The number of 16S rRNA copies per gram of sediment was evaluated by qPCR and ranged from 2.36×10^3^ to 1.7×10^6^ copies.g^-1^. Some samples had low cell numbers such as N11-1 and N11-4 with 2.59×10^4^ and 2.36×10^3^ copies.g^-1^, respectively. The sample with the highest density was N14.1 with 1.7×10^6^ copies.g^-1^ (Suppl. Figure 2 and Suppl. Table 3). No amplification occurred in sample N06-4 In the samples N11, N13 and N14, we observed a decrease in cell number with sediment depth, but this difference was not significant (Suppl. Figure 2).

## Discussion

The discovery of asphalt seeps in North São Paulo Plateau was an important milestone in studies of hydrocarbon seep environments and their associated chemosynthetic communities. This asphalt seep is similar to asphalt systems found in Campeche Knolls of southern Gulf of Mexico (MacDonald et al. 2004) and Angola Margin (Jones et al. 2014). However, Fujikura et al. (2017) analysed the oil from the asphalt seep and their results indicated that this system was not capable of sustaining chemosynthetic communities. Nevertheless, our study was the first of investigate the diversity of bacterial community using next generation sequencing in asphalt seep and non-asphalt seep sediments in the North São Paulo Plateau.

Other studies developed in deep-sea surface sediments found similar values of observed species, Chao1 and Shannon indices (Mahmoudi et al. 2014; Zhang et al. 2015), indicating that these environments harbor highly diverse microbial communities, possibly due to their temporal stability, partitioning of resources and niche diversity, allowing the coexistence of distinct microbial metabolic traits (Lozupone and Knight 2007; Zinger et al. 2011; Bienhold et al. 2016). Differences of alpha diversity values were not observed between samples at the asphalt seep and non-asphalt seep areas.

Beta-diversity analysis showed that the microbial communities distribution were not influenced by sediment depth or presence/absence of asphalt seep. Despite this, we observed a prevalence of some taxonomic groups accordingly to sediment depth. For example, four of six samples from 0-1 cm layer had as the most abundant OTU an Acidimicrobiia from Actinomarinales order, while in the second layer 1-4 cm, the most abundant OTU in three of five samples was an Alphaproteobacteria from Rhodovibrionales order (Figure 4). Jiang et al. (2018) observed that communities from surface sediments (0-4 cm) were more similar between them than communities from bottom sediments, independently whether samples were asphalt or non-asphalt seeps (16-20 cm). In our study, core sediments were sliced in two surface sediment samples (0-1 and 1-4 cm), which may explain the homogeneity between layers caused by dispersion or even by the mixture of sediments by deep-sea water currents (Meadows and Meadows 1994; Bienhold et al. 2016).

Proteobacteria and Actinobacteria comprised the prevalent phyla found in the samples, a pattern commonly observed in marine sediments throughout the world. However, at class level, we found a distinct bacterial community composition, dominated by Alphaproteobacteria, Acidimicrobiia, Gammaproteobacteria and Deltaproteobacteria, in contrast with marine sediments from other regions of the globe, where the predominant taxa in general, from most to least abundant, are Gammaproteobacteria, Deltaproteobacteria, Planctomycetes, Actinobacteria and Acidobacteria (Schauer et al. 2010; Zinger et al. 2011; Jacob et al. 2013).

The high abundance of Alphaproteobacteria and Acidimicrobiia in our samples may be explained by the presence of oil from the asphalt seep at São Paulo Plateau (Aguiar et al. 2014; Fujikura et al. 2017). Some Alphaproteobacteria taxa are able to degrade hydrocarbon such as the Rhodobacteraceae family (Kostka et al. 2011; Bacosa et al. 2018), which composed 4.5% of sequences in our samples. Bacosa et al. (2018) found an increase in the relative abundance of Rhodobacteraceae in oil treatments and, using a metagenomic approach, they could also reconstruct seven genomes, one of them classified as Rhodobacteraceae and possessing several aromatic degradation genes. We found a high abundance of the Kiloniellaceae in our samples (13%), a family which is represented by the single genera *Kiloniella* and the type species *Kiloniella laminariae* (Wiese et al. 2009; Imhoff and Wiese 2014). Wiese et al. (2009) showed by phylogenetic analysis that *Kiloniella laminariae* clustered with an uncharacterized bacterium from hydrothermal plumes and this group forms a large cluster with *Terasakiella pusilla and Thalassospira* species. Jiang et al. (2018) detected in the same area we have studied the hydrocarbon-degrading bacteria *Thalassospira xianhensis* using PCR-DGGE method.

Alphaproteobacteria contains several species which are highly abundant in superficial pelagic environments and have a broad spatial distribution. The most common example is *Pelagibacter ubique*, a ubiquitous Alphaproteobacteria present in all oceans that have important functions in biogeochemical cycles (Morris et al. 2002; Sunagawa et al. 2015). However, some studies in deeper pelagic environments also found Alphaproteobacteria composing most of the microbial community (Konstantinidis et al. 2009; Eloe et al. 2011). Therefore, this proximity between deep seawater and sediment surface may allow microbial community interchange, since both environments have similar chemical variables and suitable habitats for these microbial populations (Hamdan et al. 2013; Walsh et al. 2016).

Acidimicrobiia was highly abundant in North São Paulo Plateau sediments. The taxon Acidimicrobiia was recently created by updating taxonomic classification of Actinobacteria phylum to include the Acidimicrobidae subclass, assigned as Acidimicrobiia class (Zhi et al. 2009). Most of our sequences assigned to this phylum belonged to the order Actinomarinales, and the two most representative OTUs were classified as uncultured actinobacterium (previously classified as OM1 clade). The OM1 clade group was also found in deep-sea waters (Eloe et al. 2011; Quaiser et al. 2011), as an important component of deep-sea sediment core microbiome in several oceans (Bienhold et al. 2016).

It is generally assumed that microbial cell densities in deep-sea sediment tend to decrease with increasing sediment depth (Orcutt et al. 2011). In our study this tendency was not observed, with differences in bacterial densities not being significant, probably by the microbial communities interchanges between deep seawater and sediment surface. Microbial densities in North Sao Paulo Plateau might vary in deeper sediment layers (> 4 cm) not achieved in our study. In addition, the low abundance of 16S rRNA gene copies corroborates with similar deep-sea sediments habitats (Jorgensen et al. 2012), indicating that these habitats in the North Sao Paulo Plateau are oligotrophic, and sustain a low abundant, but diverse microbial community.

## Conclusions

Bacterial communities in the North Sao Paulo Plateau are diverse, despite their low abundance, and are dominated by the classes Alphaproteobacteria and Acidimicrobiia. This community structure differs from other communities from similar environments, in which Gammaproteobacteria are usually more abundant. We also found a high number of unclassified sequences mainly related to Actinomarinales order, suggesting that this environment can harbor groups poorly explored to date. The dominance of Alphaproteobacteria potentially involved with hydrocarbon degrading might be likely related to the presence of asphalt seeps, however further studies are needed to answer this question.

## Acknowledgements

We would like to thank Japan Agency for Marine-Earth Science and Technology (JAMSTEC), the Oceanographic Institute of the São Paulo University (IOUSP), the Brazilian Geological Survey (CPRM), Petróleo Brasileiro S.A. (Petrobras) and the Embassy of Japan in Brazil for assistance in this study. We would also like to thank Fundação de Amparo a Pesquisa do Estado de São Paulo (FAPESP) for financial support (Project number: 2013/09159-2) and CNPq for scholarship provided to A.O.S.L (Process 311010/2015-6); the operating team of the HOV Shinkai 6500 and the crew of the R/V Yokosuka for assistance with the survey; and all team of Laboratório de Ecologia Microbiana (LECOM) for productive discussions about our methods and results, and Kleber do Espirito-Santo Filho for help with maps.

## Data

The nucleotide sequence data reported are available in the NCBI under BioProject PRJNA562874.

## Authorship

The author LQ, RD, CN, PS, AL, YN, KF, HK and VP designed study, LQ, RD,DG,AS and VP performed research, LQ, AB, RD and DG analysed data; LQ, AB, RD and DG contributed new methods or models; and LQ, AB, RD, CN and VP wrote the paper.

