## Supplementary material for "Bacterial diversity in deep-sea sediments under influence of asphalt seep at the São Paulo Plateau"

Supplementary figure 1. Principal Coordinates Analysis (PCoA) of seven sediment samples from North São Paulo Plateau considering asphalt seep influence and deep layer for Bray-Curtis, Jaccard, Unweighted and Weighted UniFrac dissimilarity matrix.

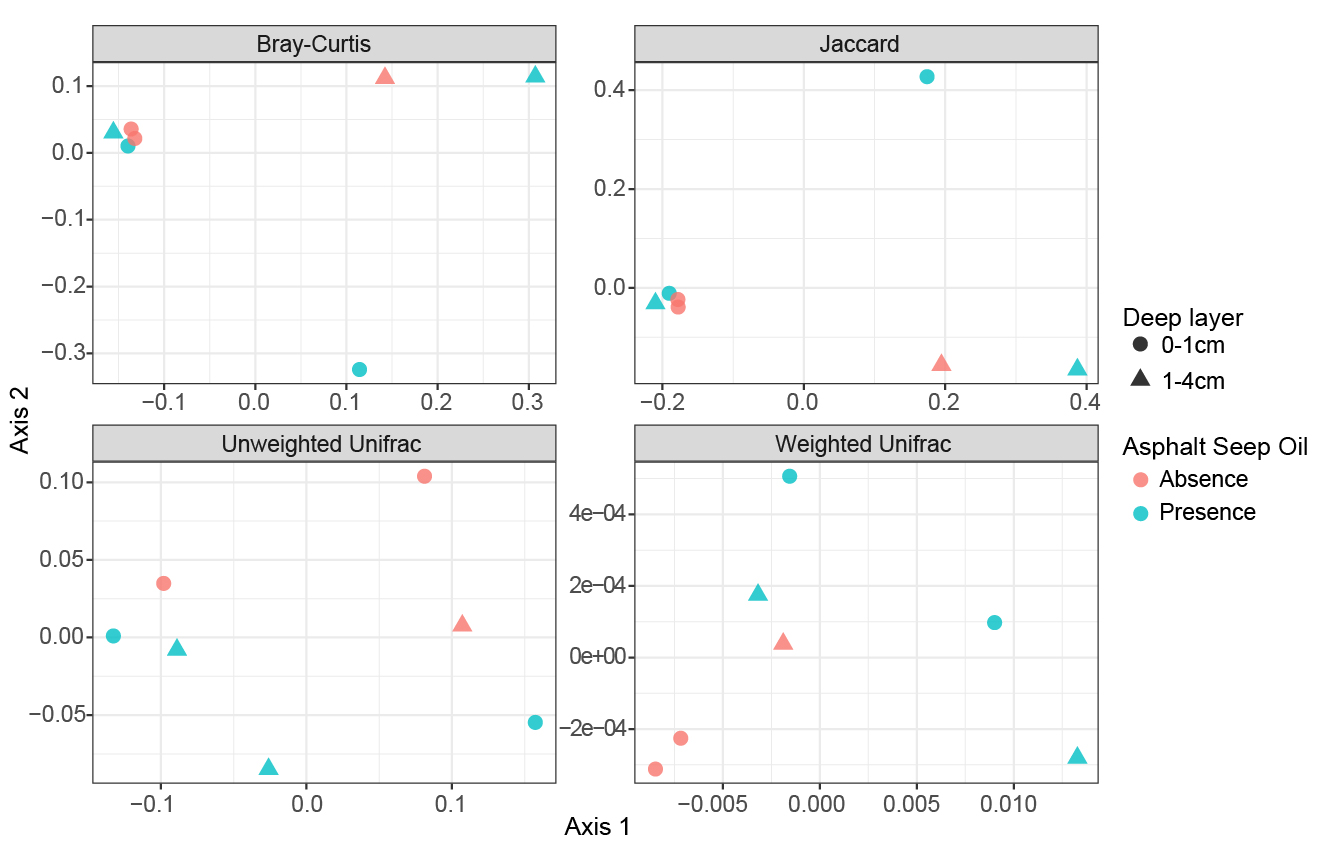

Supplementary figure 2. Bacterial 16S rRNA number of copies estimated for each sample and layer. Sediment layers were compared by paired Wilcoxon-Mann Whitney test.

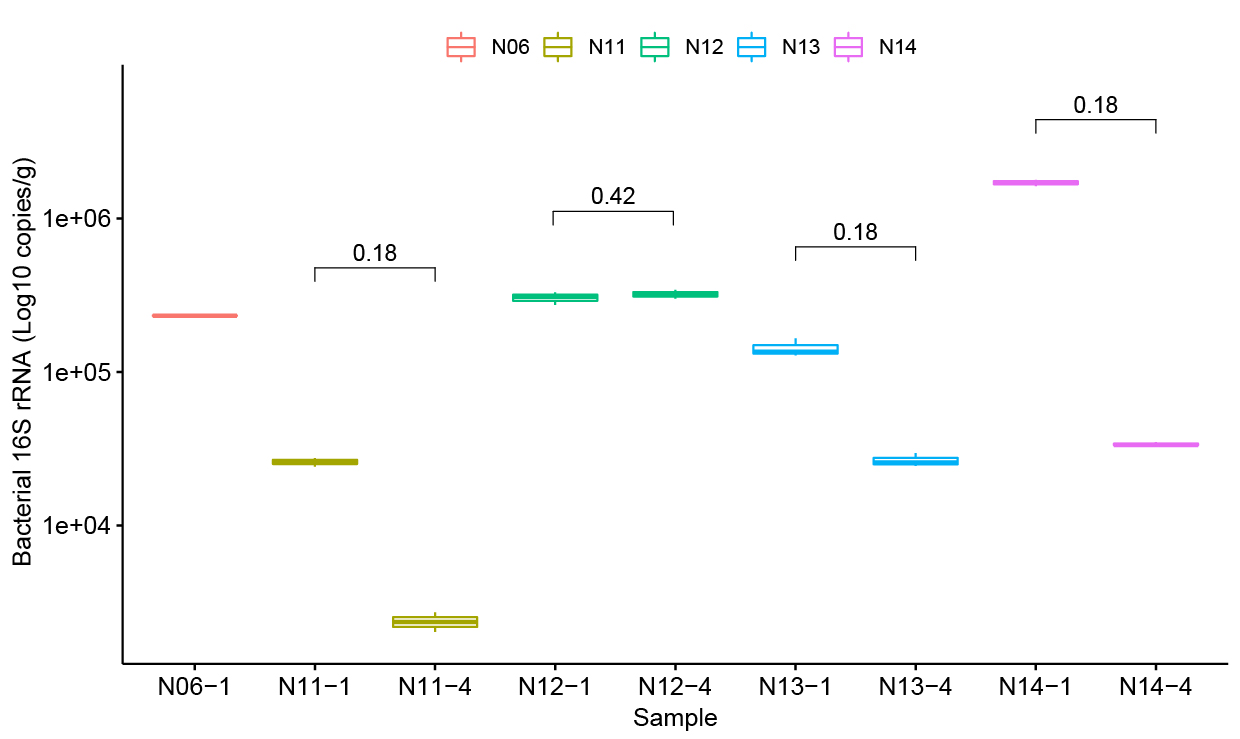

Supplementary table 1. Differences of alpha-diversity between asphalt and non-asphalt seep areas. Means were compared using *t-*test.

| Alpha-diversity index | Asphalt seep | | |
| --- | --- | --- | --- |
|  | Absent | Present | P value |
| Observed | 1878 | 1768 | 0.7628 |
| Shannon | 8.021 | 8.092 | 0.5227 |
| Simpson | 0.983 | 0.986 | 0.0243* |
| Chao1 | 2363 | 2252 | 0.8479 |

Supplementary table 2. Results of PERMANOVA analysis of sediment samples from North São Paulo Plateau considering asphalt seep influence and deep layer for using Bray-Curtis, Jaccard, Unweighted and Weighted Unifrac dissimilarity matrix.

| Variable | Matrix distance | F | R2 | P |
| --- | --- | --- | --- | --- |
| Oil | Bray-Curtis | 0.697 | 0.125 | 0.664 |
|  | Jaccard | 0.766 | 0.134 | 0.626 |
|  | Weighted Unifrac | 2.796 | 0.451 | 0.217 |
|  | Unweighted Unifrac | 0.848 | 0.172 | 0.450 |
| Layer | Bray-Curtis | 1.230 | 0.221 | 0.336 |
|  | Jaccard | 1.162 | 0.204 | 0.366 |
|  | Weighted Unifrac | 0.260 | 0.042 | 0.587 |
|  | Unweighted Unifrac | 0.473 | 0.096 | 0.708 |

Supplementary table 3. Cell number of Bacteria 16S rRNA gene in samples of North São Paulo Plateau. SD: standard deviation.

|  | Samples | | | | | |
| --- | --- | --- | --- | --- | --- | --- |
| Layers | N06 | SD | N11 | SD | N12 | SD |
| 1 cmbsf | 2.32×10^5^ | 1.39×10^3^ | 2.59×10^4^ | 1.64×10^3^ | 3.03×10^5^ | 2.85×10^4^ |
| 4 cmbsf | - | - | 2.36×10^3^ | 3.45×10^2^ | 3.21×10^5^ | 2.13×10^4^ |
| Layers | N13 | SD | N14 | SD |  |  |
| 1 cmbsf | 1.42×10^5^ | 1.96×10^4^ | 1.70×10^6^ | 7.73×10^4^ |  |  |
| 4 cmbsf | 2.65×10^4^ | 2.66×10^3^ | 3.38×10^4^ | 9.04×10^2^ |  |  |

Obs.: Sample N06-4cm did not amplify.
